# Human serum supplementation promotes *Streptococcus mitis* growth and induces specific transcriptomic responses

**DOI:** 10.1101/2022.12.13.520276

**Authors:** Yahan Wei, Camille I. Sturges, Kelli L. Palmer

## Abstract

*Streptococcus mitis* is a normal member of the human oral microbiota and a leading opportunistic pathogen causing infective endocarditis (IE). Despite the complex interactions between *S. mitis* and human host, understanding of *S. mitis* physiology, as well as its mechanisms of adaptation to host-associated environments, is inadequate, especially when compared with other IE bacterial pathogens. This study reports growth-promoting effects of human serum on *S. mitis* and other pathogenic streptococci, including *S. oralis, S. pneumoniae*, and *S. agalactiae*. Using transcriptomic analyses we identified that, with the addition of human serum, *S. mitis* down-regulates uptake systems for metal ions and sugars, fatty acid biosynthetic genes, and genes involved in stress response and other processes related with growth and replication. *S. mitis* up-regulates uptake systems for amino acids and short peptides in response to human serum. Zinc availability and environmental signals sensed by the induced short peptide binding proteins were not sufficient to confer the seen growth-promoting effects. More investigation is required to establish the mechanism for growth promotion. Overall, our study contributes to the fundamental understanding of *S. mitis* physiology under host-associated conditions.

**Significance:** *S. mitis* is exposed to human serum components during commensalism in the human mouth and bloodstream pathogenesis. However, the physiological effects of serum components on this bacterium remain unclear. Using transcriptomic analyses, *S. mitis* biological processes that respond to the presence of human serum were revealed, providing the fundamental understanding of *S. mitis* adaptations towards human host conditions.

## Introduction

*Streptococcus mitis* is a member of the mitis group streptococci (MGS) and a pioneer human oral colonizer (1, 2). Through invasive dental procedures (e.g. tooth extraction and scaling) as well as daily dental hygiene activities (e.g. toothbrushing and flossing), *S. mitis* gains access to the bloodstream and causes transient bacteremia, imposing a health risk to people with compromised immunity (3). Indeed, *S. mitis* is one of the leading causes of bacteremia and infective endocarditis (IE), which present a mortality rate of ∼ 18 to 40% (4–6). Conversely, *S. mitis* also has beneficial functions, as it can produce antimicrobials that inhibit growth of human gingivitis pathogens (7, 8), and its lysate can activate the aryl hydrocarbon receptor (AhR) of oral epithelial cells, leading to wound healing (9). Moreover, decreased *S. mitis* in the oral microbiota is a biomarker for pancreatic cancer (10). However, the specific molecular mechanisms underlying interactions between *S. mitis* and the human host remain largely unknown. Furthermore, *S. mitis* physiology, compared to other bacteremia and IE bacterial pathogens (e.g. *Staphylococcus aureus* and enterococci), is poorly understood.

The current knowledge of *S. mitis* physiology as a human commensal and opportunistic pathogen is largely adapted from studies in another MGS, *S. pneumoniae* (11). *S. pneumoniae* is a significant human pathogen that causes ∼1 million annual deaths of children < 5 years old worldwide and shares > 99% 16S rRNA sequence identity with *S. mitis* (12–14). Many basic physiological processes and several cell surface components are conserved between *S. mitis* and *S. pneumoniae* (15–17). For example, *S. mitis* encodes 67-82% of known pneumococcal virulence factors (18). Intranasal immunization with *S. mitis* provides protection against *S. pneumoniae* (19), suggesting similarities in both cell surface structures and interactions with host immunity between these bacteria. However, whole chromosomal DNA identity between *S. mitis* and *S. pneumoniae* is <60% (14). Genes unique to *S. mitis* are involved in processes including oligopeptide binding, amino acid metabolism, gene regulation, and virulence (17). Such genetic differences possibly contribute to the different niches these bacteria colonize in the human host and the diseases caused by each. A comprehensive understanding of *S. mitis* physiology, particularly in conditions that mimic human host environments encountered by this bacterium during colonization and infection, is fundamental to an understanding of its interactions with the human host.

Transcriptomic analyses provide a thorough picture of the physiological status of an organism and have been widely used in bacterial studies (20), yet are rarely applied to *S. mitis*. Thus far, only a few transcriptomic studies have been conducted in *S. mitis*, each designed to resolve 1) responses to competence stimulating peptides (CSPs) (21), 2) interspecies communication with *S. pneumoniae* via secreted short peptides (22), and 3) genetic differences against other major MGS species cultured in rich, undefined laboratory medium (23). Here, we assessed the growth and transcriptional responses of *S. mitis* to human serum. When colonizing the gingival pocket in the human oral cavity, *S. mitis* is exposed to gingival crevicular fluid, which contains serum transudate (24). When colonizing the bloodstream, *S. mitis* is also constantly exposed to serum components. Thus, adaptation to human serum components is likely critical for this bacterium to colonize these environments. In this study, we observed growth-promoting effects of human serum on several streptococcal species (including *S. mitis, S. oralis, S. pneumoniae*, and *S. agalactiae*), but not bacteria from other genera. Transcriptomic analyses were performed to reveal the responses of *S. mitis* to the addition of human serum, whose presence alleviates stresses and provides key nutrients, providing information to understand *S. mitis* adaptation to host-associated environments.

## Results and discussion

### Addition of human serum promotes the growth of some streptococci

To test how human serum addition affects bacterial growth, selected bacterial species were grown in either chemically defined medium (CDM) or CDM supplemented with complete human serum to a final concentration of 5% (v/v). Significant increases in cell yield as measured by colony forming units (CFUs) were observed for *S. mitis, S. oralis, S. pneumoniae*, and *S. agalactiae* cultured with human serum. This was not observed for *S. pyogenes* and bacteria from other genera (*S. aureus, Enterococcus faecalis*, and *Escherichia coli*) (Fig. 1A). Additional *S. mitis* and *S. oralis* strains were also tested. These strains were reported recently (25) and were isolated from the oral cavities of volunteers (Table 3, oral isolates) or from the bloodstream of hospitalized patients (Table 3, blood isolates). Human serum significantly promoted growth of some, but not all, of the isolates (Fig 1B).

**Table 1:**
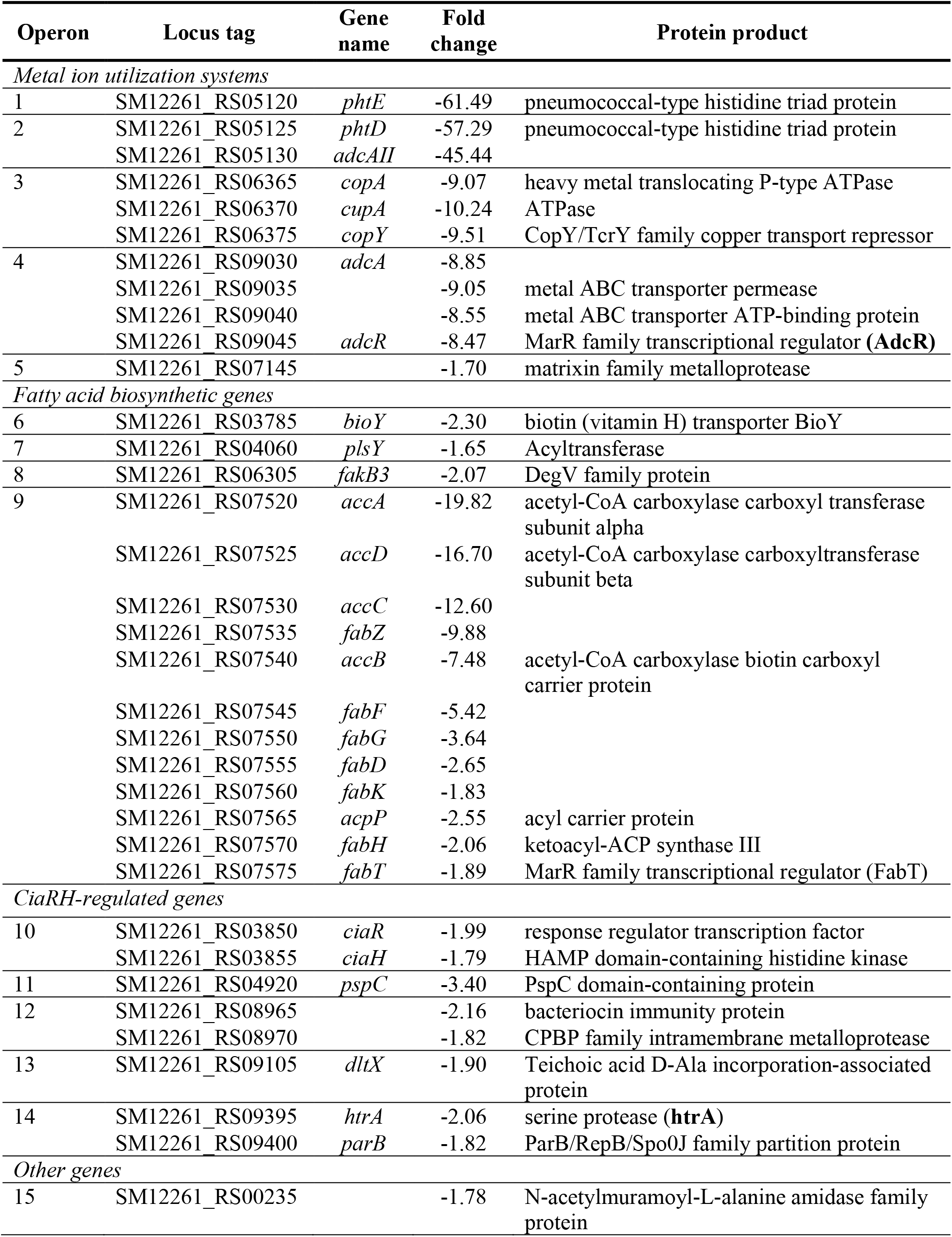
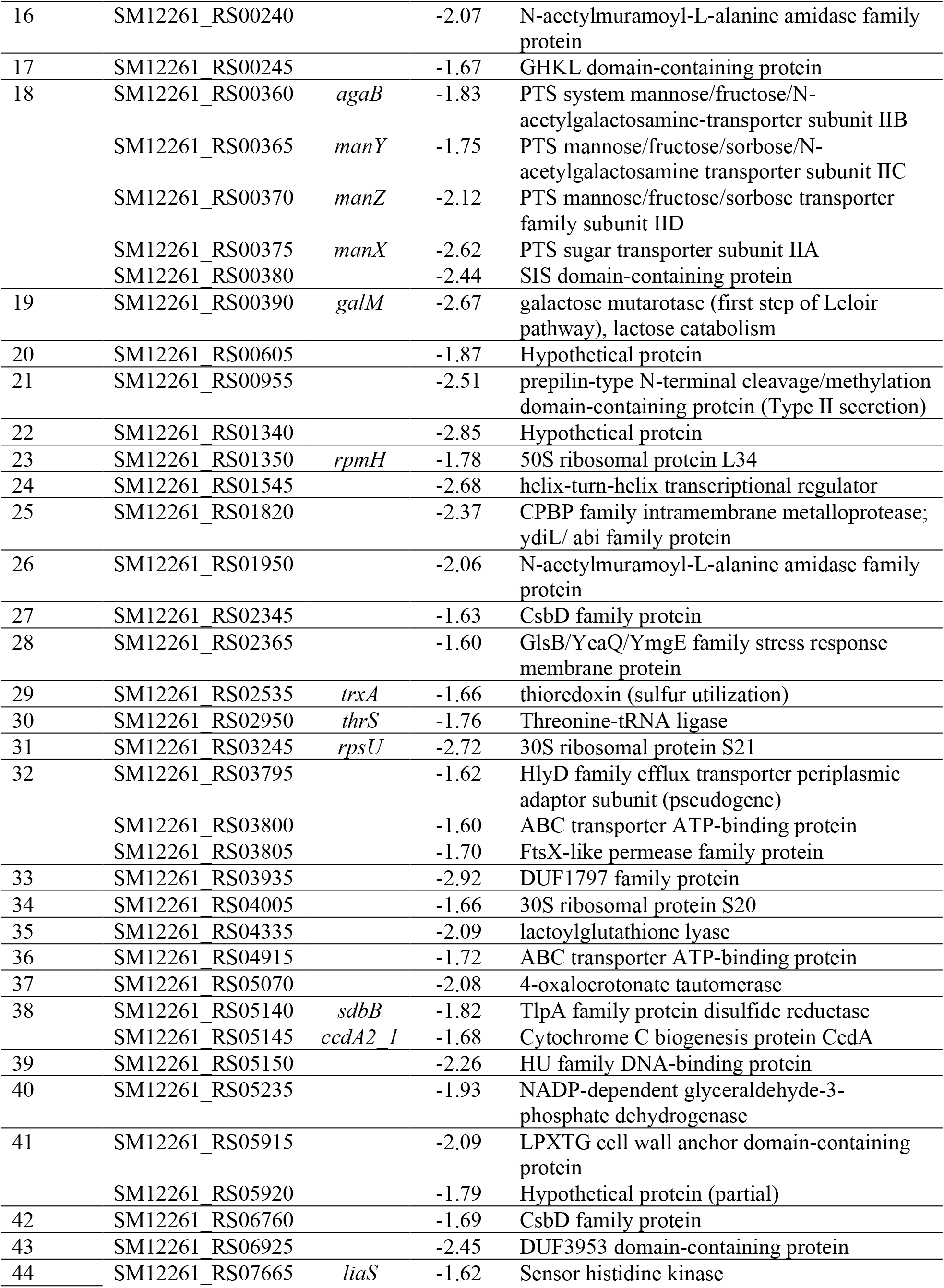
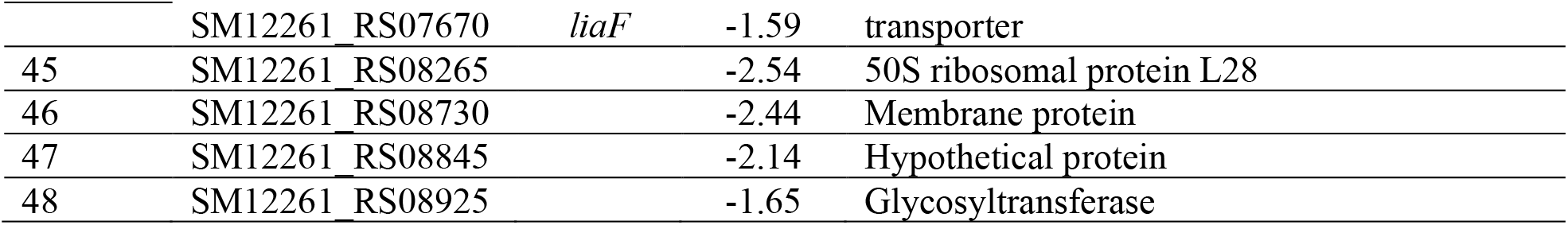
Genes downregulated upon the addition of human serum

**Table 2:** Genes upregulated upon the addition of human serum

| Operon | Name | Gene | Fold change | Protein product |
| --- | --- | --- | --- | --- |
| *CiaRH-regulated gene* |  |  |  |  |
| 1 | SM12261_RS01270 | *manL* | 1.69 | Mannose-specific IIB subunit of PTS system |
| *Metalloenzymes* |  |  |  |  |
| 2 | SM12261_RS00935 |  | 4.15 | zinc-dependent alcohol dehydrogenase family protein |
| 3 | SM12261_RS01275 | *adhP* | 4.99 | alcohol dehydrogenase |
| 4 | SM12261_RS01285 |  | 1.54 | NCS2 family permease (nucleobase transporter) |
| 5 | SM12261_RS06110 |  | 1.51 | FAD: protein FMN transferase |
|  | SM12261_RS06115 |  | 1.51 | NAD(P)H-dependent oxidoreductase |
| 6 | SM12261_RS09120 | *glpO* | 1.99 | Type I glycerol-3-phosphate oxidase |
| *Amino acid transportation* |  |  |  |  |
| 7 | SM12261_RS01690 | *glnA* | 1.95 | Type I glutamate--ammonia ligase |
| 8 | SM12261_RS05465 |  | 1.98 | ABC transporter permease subunit |
|  | SM12261_RS05470 |  | 1.80 | amino acid ABC transporter ATP-binding protein |
| 9 | SM12261_RS07175 |  | 3.32 | amino acid ABC transporter ATP-binding protein |
|  | SM12261_RS07180 |  | 3.84 | transporter substrate-binding domain-containing protein |
|  | SM12261_RS07185 |  | 3.32 | amino acid ABC transporter permease |
|  | SM12261_RS07190 |  | 2.83 | amino acid ABC transporter permease |
| *Peptide uptake* |  |  |  |  |
| 10 | SM12261_RS06745 | *aliB* | 1.70 | peptide ABC transporter substrate-binding protein; *amiA aliA/B* homolog |
| 11 | SM12261_RS07850 |  | 1.75 | peptide ABC transporter substrate-binding protein (AliB-like ORF2) |
| *Hypothetical proteins* |  |  |  |  |
| 12 | SM12261_RS02965 |  | 1.85 | ABC transporter ATP-binding protein |
| 13 | SM12261_RS08945 |  | 1.76 | DUF4097 family beta strand repeat protein |
|  | SM12261_RS08950 |  | 1.76 | Hypothetical protein |

**Table 3:** Strains and plasmids used in this research

| Name | Description | Reference |
| --- | --- | --- |
| **Bacterial strains** |  |  |
| *Escherichia coli* K-12<br>MG1655 | Model *E. coli* strain | (70) |
| *Staphylococcus aureus*<br>RN4220 | Laboratory strain of *S. aureus*, a mutant strain derived from clinical isolate NCTC8325 | (71, 72) |
| *Enterococcus faecalis* |  |  |
| OG1RF | Rifampin- and fusidic acid-resistant derivative of a human oral cavity isolate | (73) |
| Merz96 | Vancomycin resistant strain isolated from human blood | (74) |
| *Streptococcus* |  |  |
| *S. mitis* ATCC 49456<sup>T</sup><br>(SM61) | Type strain of *S. mitis* | ATCC |
| *S. mitis* ΔaliB | SM61 with coding region of SM12261_RS06745 replaced with *ermB* | This study |
| *S. mitis* O1-2 | Oral isolate obtained from healthy individual | (25) |
| *S. mitis* O7-1 | Oral isolate obtained from healthy individual | (25) |
| *Streptococcus* sp. 1643 | Mitis group *Streptococcus* isolated from infective endocarditis patient | (75) |
| *S. oralis* ATCC 35037<sup>T</sup> | Type strain of *S. oralis* | ATCC |
| *S. oralis* O3-1 | Oral isolate obtained from healthy individual | (25) |
| *S. oralis* O8-3 | Oral isolate obtained from healthy individual | (25) |
| *S. oralis* S33 | Strain isolated from clinical blood sample | (25) |
| *S. oralis* S50 | Strain isolated from clinical blood sample | (25) |
| *S. oralis* S51 | Strain isolated from clinical blood sample | (25) |
| *S. pneumoniae* D39 | Clinically isolated strain, serotype 2 | (27) |
| *S. agalactiae* COH1 | Clinically isolated strain, type III capsular polysaccharide | (76) |
| *S. pyogenes* NZ131 | Clinically isolated strain, serotype M49 | ATCC |
| **Plasmids** |  |  |
| pMSP3535 | Plasmid used to obtain *ermB* gene | (77) |

**Fig 1:**
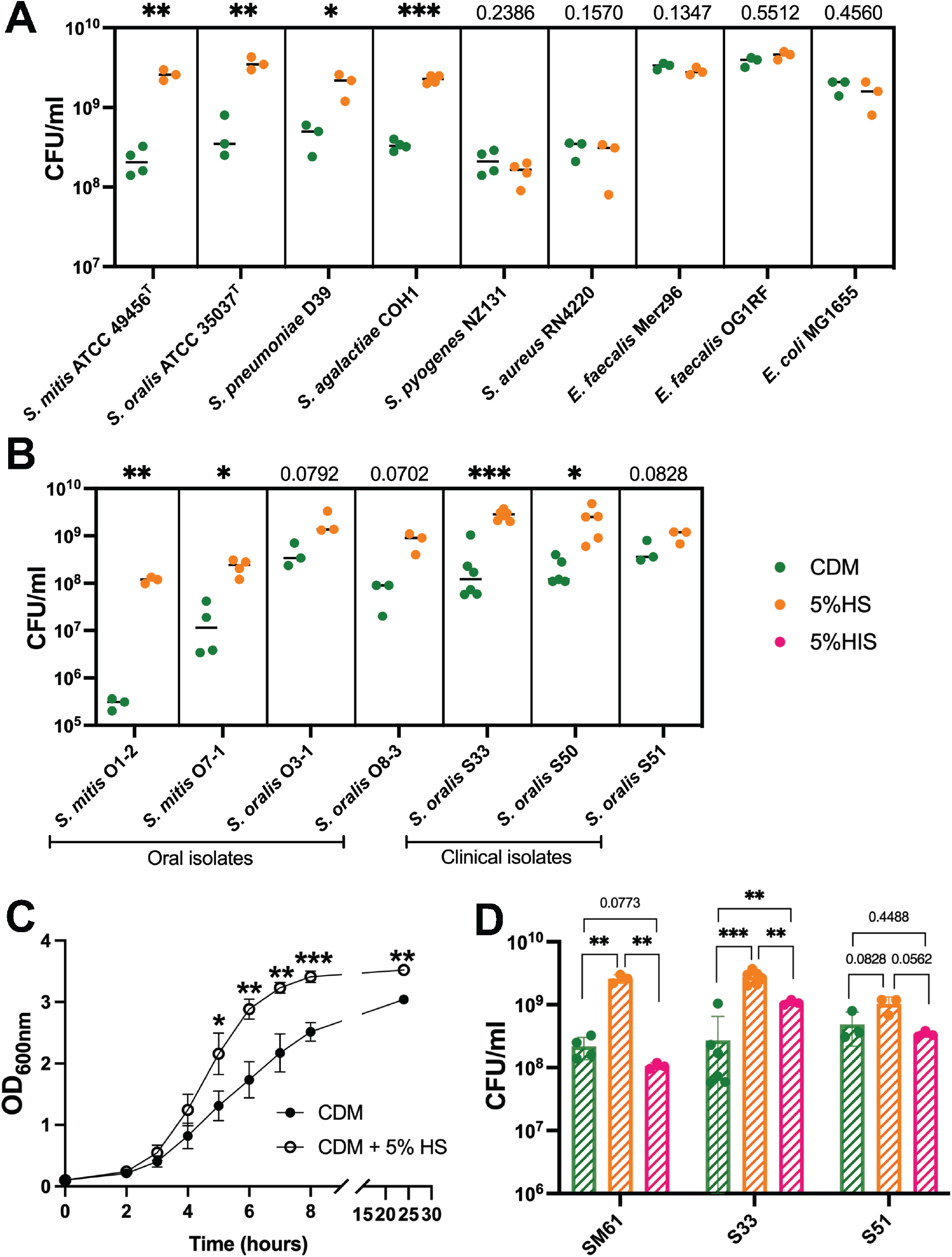
Serum supplement promotes the growth of some streptococci. A. Cell concentrations (CFU/ml) of the indicated bacteria grown to stationary phase in either chemically defined medium (CDM, green dots) or CDM with 5% (v/v) supplement of complete human serum (HS, orange). B. CFU/ml of the indicated MGS isolates grown to stationary phase in either CDM (green dots) or HS (orange dots). C. Growth curve of *S. mitis* ATCC 49456^T^ (SM61) grown in either CDM or CDM with 5% (v/v) supplement of HS. D. Stationary phase cell densities (CFU/ml) of SM61, *S. oralis* S33 (S33), and *S. oralis* S51 (S51) grown in CDM (green), HS (orange), or CDM with 5% (v/v) supplement of heat-inactivated human serum (HIS, pink). At least three biologically independent replicates were obtained for each tested condition. Statistical analyses were performed with two-tailed *T*-test, with *F*-test conducted to verify whether compared groups have equal variances. Significant difference is determined by *P*-value < 0.05. Nonsignificant *P*-values were labeled within graph; “*” indicates 0.05 > *P*-value > 0.01; “**” indicates 0.01 > *P*-value > 0.001; “***” indicates 0.001 > *P*-value.

For *S. mitis* ATCC 49456^T^ (SM61), addition of human serum resulted in a 10-fold average increase of cell yield at stationary phase (Fig. 1A) and significantly decreased generation time from 46.8 (±1.6) min in CDM to 39.1 (±0.8) min in serum-supplemented CDM with a T-test *P*-value of 0.002 (Fig. 1C). Interestingly, when supplemented with heat-inactivated serum, growth of SM61 and the clinical isolate *S. oralis* S33 decreased significantly compared to those grown with untreated serum (Fig 1D). Another clinical isolate, *S. oralis* S51, had no significant differences in growth with either heat-inactivated or untreated serum (Fig 1D).

Overall, these data suggest that temperature-sensitive component(s) is/are involved in the growth promoting effects of human serum, and that the effects of human serum on streptococcal growth are species- and strain-dependent.

### Transcriptomic analysis of the biological processes involved in growth promotion with serum addition

To analyze the transcriptional responses of SM61to human serum addition, RNA sequencing was performed. For these experiments, *S. mitis* cells in exponential phase (∼0.3 OD_600nm_) exposed to 5% human serum for 30 minutes were compared with control, unexposed cultures. This approach was taken to minimize the confounding effects of different growth rates/yields between the two conditions on transcriptional analysis, and to focus on initial responses of *S. mitis* to human serum. Expression values were represented by reads per kilobase of transcript, per million mapped reads (RPKM). Genes differentially regulated in response to serum were determined by ≥1.5-fold absolute changes at the transcript level, corresponding false discovery rate (FDR) *P*-value <0.01, and maximum RPKM >10. By these parameters, a total of 77 downregulated genes (Table 1) and 19 upregulated genes (Table 2) were obtained, which, according to previous genomic annotations (17, 26), are all shared between *S. mitis* and *S. pneumoniae*. According to the annotated *S. mitis* locus structures and the transcription start sites of their homologs in *S. pneumoniae* (27), these down- and up-regulated genes belong to approximately 48 and 13 operons, respectively (Table 1 & 2). Transcriptional changes for a subset of genes were confirmed via quantitative reverse transcriptase PCR (qRT-PCR) (Fig. 2).

**Fig 2:**
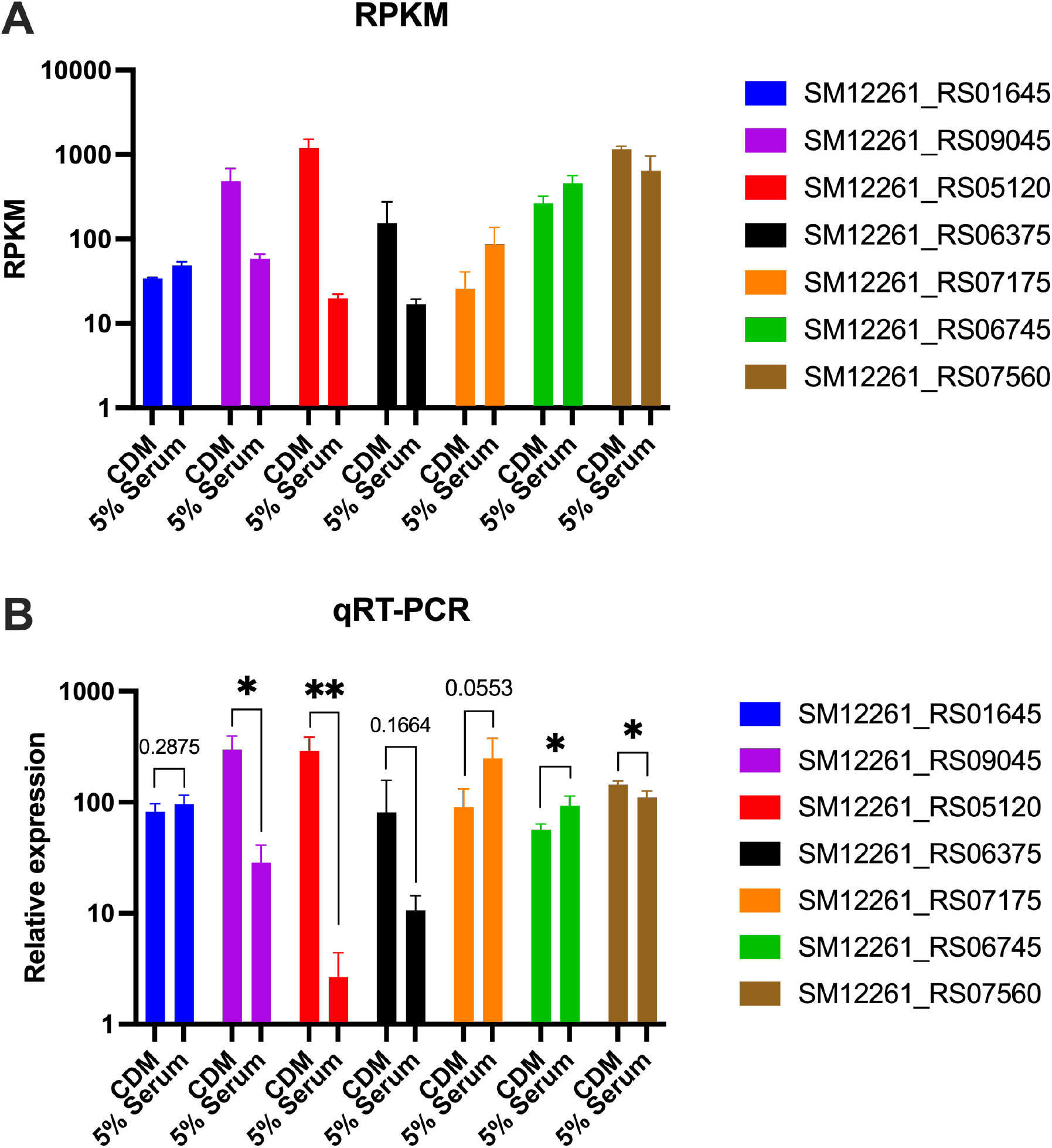
Relative expression levels of selected SM61 genes detected through RNA sequencing (A) and qRT-PCR (B). Except SM12261_RS01645 (encodes an iron utilization gene), all other genes are differentially expressed in RNA sequencing analyses. A. RPKMs of each selected gene. B. transcript levels of each selected gene were normalized to that of 16S rRNA from the same sample with the ΔΔCt method. For each tested condition, three biologically independent samples were obtained. Statistical analyses were performed with two-tailed T-test, with *F*-test conducted to verify whether compared groups have equal variances or not. Significant difference determined by *P*-value < 0.05. Nonsignificant *P*-values were labeled within graph; “*” indicates 0.01 < *P*-value < 0.05; “**” indicate 0.001 < *P*-value < 0.01; “***” indicate 0.0001 < *P*-value < 0.001.

In general, the regulated genes are involved in biological processes of 1) transportation of metal ions (zinc and copper), sugars, amino acids, and short peptides, 2) fatty acid biosynthesis, including type II fatty acid synthetic (FASII) genes, an acyltransferase, and an exogenous fatty acid binding protein, 3) putative stress sensing regulators and stress response genes, and 4) proteins with unknown functions (Table 1 & 2).

### Downregulation of metal ion transportation

The most downregulated genes in *S. mitis* SM61 supplemented with serum are *phtE, phtD*, and *adcII*, which are associated with Zn utilization (Table 1). Other Zn utilization genes of the *adcR* locus are also downregulated > 5-fold. While zinc is present in human serum at an estimated concentration of 9 – 17 μM (28, 29), CDM does not contain added Zn. Thus, addition of serum increases the amount of available Zn. *S. mitis* likely downregulates Zn uptake genes after serum addition to maintain metal ion homeostasis. However, addition of Zn into CDM does not result into accelerated growth, nor increase the stationary phase OD_600nm_ values of *S. mitis* (Fig. S1). Therefore, increased Zn availability alone does not explain the growth promoting effects of human serum.

In *S. pneumoniae*, the cellular concentration of Zn affects the expression of Zn uptake genes and those involved in Zn export (*sczD*) (30), Mn uptake (*psaABC*) (31), and Cu export (*copA*) (32) through modulating activities of their corresponding transcriptional regulators. Among these genes, homologs of the *copY*-*cupA*-*copA* operon that encodes a transcriptional repressor (CopY), Cu chaperone (CupA), and Cu exporter (CopA) are downregulated in *S. mitis* upon serum addition. In the presence of Zn, pneumococcal CopY binds DNA and represses transcription of its target genes (32); thus, the observed downregulation of the *copY*-*cupA*-*copA* likely results from increased Zn abundance due to serum addition. Interestingly, other genes that are expected to be regulated in response to increased Zn availability were not differentially expressed in our data set (Table S1), suggesting a hierarchical regulatory relationship among these metal utilization systems in *S. mitis*.

### Downregulation of fatty acid biosynthetic genes

Aside from the Zn utilization genes, genes encoding the acetyl-CoA carboxylase complex (*accADC*) are the next most downregulated genes in this data set for fold changes >10 (Table 1). These genes belong to the type II fatty acid biosynthetic (FASII) pathway for *de novo* fatty acid biosynthesis. In *S. pneumoniae*, incorporation of exogenous lipids downregulates the expression of FASII genes through the activities of transcriptional regulator FabT (33). Human serum is a rich source of lipids (34). Evidence that streptococci utilize serum fatty acids already exists (35, 36); thus, downregulation of the FASII genes in *S. mitis* with the addition of human serum is intuitive. Indeed, all FASII genes are downregulated in *S. mitis* after serum supplementation (Table 1).

Interestingly, downregulation of the biotin transporter gene *bioY* (SM12261_RS03785), a homolog of the exogenous fatty acid binding protein FakB3 (SM12261_RS06305), and acyltransferase gene *plsY* (SM12261_RS04060) were also observed. Biotin is a cofactor for FASII carboxylation enzymes (37). Thus, it is possible that expression of *bioY* correlates with that of the FASII genes, through an as yet unknown regulatory mechanism in *S. mitis*. Both *fakB3* and *plsY* are involved in fatty acid utilization. Specifically, exogenous fatty acid is recognized and bound by a cell surface protein (FakB), phosphorylated by a fatty acid kinase (FakA), and then used for synthesis of phosphatidic acids by acyltransferases that include PlsY (38). *S. mitis* encodes three copies of *fakB*, SM12261_RS06970, SM12261_RS05155, and SM12261_RS06305, corresponding to homologs of the pneumococcal fatty acid binding protein FakB1, FakB2, and FakB3. In *S. pneumoniae*, these FakB proteins recognize fatty acids that differ by their acyl chain length and saturation status (39). Exact functions of *S. mitis* FakB proteins still await further studies, as do the transcriptional regulatory mechanisms that control the expression of *fakB* genes. Interestingly, among the three *fakB* genes, only *fakB3* was differentially expressed after serum addition. Similarly, among the three known acyltransferases in *S. mitis* (*plsC* SM12261_RS03010, *plsY*, and *plsX* SM12261_RS00220), only *plsY* was differentially expressed after serum addition. Such differences in expression status suggest different functions of these genes and/or different regulatory systems that individually fine-tune the expression of these genes in response to environmental changes.

### Downregulation of stress sensing regulators and stress response genes

Several genes with activities in transcriptional regulation and stress response are downregulated in response to human serum, suggesting that serum addition alleviates some stresses, potentially triggering growth-promoting effects. Among these regulatory systems, homologs of the two-component system (2CS), CiaRH (*ciaR*: SM12261_RS03850, *ciaH*: SM12261_RS03855), and *liaFS* (*liaF*: SM12261_RS07670, *liaS*: SM12261_RS07665), which are part of the three-component system (3CS) LiaFSR, are downregulated, as well as genes that are predicted to be regulated by these systems. LiaFSR is conserved among several Gram-positive bacteria (40–43), senses structural and compositional changes in the bacterial membrane and cell-wall components, and mediates stress responses (44, 45). Thus, the downregulation of *liaFS* possibly results from altered lipid compositions caused by the addition of serum, as indicated by the downregulation of FASII genes (Table 1). Among the known operons that are directly regulated by LiaR (46), besides the *liaFS* genes, only the *pspC* homolog was differentially expressed, which is also regulated by CiaR.

CiaRH is a conserved 2CS shared among many pathogenic streptococcal species, including *S. pneumoniae, S. agalactiae*, and *S. pyogenes* (47). It mediates resistance to stresses such as oxidative stress, acid, and antibiotics (48) and controls several growth and virulence-related processes such as cell wall biosynthesis and biofilm formation (15, 46, 47, 49). In response to unknown environmental stimuli, membrane-embedded histidine kinase CiaH phosphorylates transcriptional regulator CiaR, which activates its DNA binding ability (50). Additionally, acetyl phosphate and other histidine kinases can also activate CiaR (50). In *S. pneumoniae*, CiaR directly regulates 18 promoters (46). Among those regulated genes, homologs to the *htrA*-*parB* operon for replication partition (SM12261_RS09395-400), bacteriocin production (SM12261_RS08965-70), cell-wall D-alanine modification *dltX* (SM12261_RS09105), virulence gene *pspC* (SM12261_RS04920), and the first gene of a mannose transporter operon (SM12261_RS01270), were all differentially regulated in response to human serum (Table 1 & 2). Six of the 18 CiaR-regulated promoters in *S. pneumoniae* drive expression of non-coding RNAs (ncRNAs) (46); however, the profile of *S. mitis* ncRNAs is still unclear, thus their expression levels were not included in this analysis.

### Genes upregulated in the presence of human serum

Nineteen genes were upregulated by *S. mitis* after the addition of human serum (Table 2). These genes include a sugar uptake gene putatively regulated by CiaRH, 6 metalloenzyme genes possibly regulated in response to changes in metal ion availability, 7 amino acid transportation genes, 2 short peptide uptake genes, and 3 hypothetical genes of unknown functions.

Glutamine synthetase GlnA is involved in regulating bacterial nitrogen metabolism, including glutamine uptake (51). Additionally, expression of glutamine transporters is co-regulated by transcriptional regulators, including CodY and GlnR (51, 52), which were not differentially regulated in our dataset. Bioinformatic analyses identified at least six glutamine transporters in *S. pneumoniae* (53). *S. mitis* encodes homologs of each of the six systems (Table S2), only one of which (SM12261_RS07175-90) is differentially regulated with serum addition. However, qRT-PCR did not confirm the statistical significance of the upregulated transcription of SM12261_RS07175 (Fig 2B). Further studies are needed to clarify the detailed nitrogen metabolic processes in *S. mitis*.

Streptococci can uptake short peptides as environmental signals and correspondingly modulate cellular processes (54). One such example is the induction of natural competence in mitis group streptococci by CSP (55). Environmental short peptides are imported through oligopeptide permeases, with different peptide-binding proteins recognizing peptides of different sequences (56). *S. mitis* encodes at least 7 different peptide-binding proteins (Table S1), among which, SM12261_RS06745 (homolog to *S. pneumoniae aliB*) and SM12261_RS07850 (*aliB*-like ORF 2, referred to hereafter as ORF 2) are significantly upregulated after serum addition. In *S. pneumoniae*, addition of peptide ligands of AliB or ORF 2 to culture media can boost bacterial growth (57, 58). To verify whether AliB is responsible for the growth-promoting effect of human serum, a Δ*aliB* deletion mutant was generated in *S. mitis*. However, no significant growth difference was observed between WT and *ΔaliB* SM61 that were grown in either plain CDM or serum supplemented CDM (Fig. 3), and the growth promoting effect of serum was still observed for *ΔaliB* (Fig. 3). Considering that deletion of the ORF2 gene in *S. pneumoniae* also failed to abolish the growth-promoting effect of the corresponding peptide ligand (58), it is possible that these peptide receptors have redundant functions in ligand binding. Additionally, human serum, as a complex nutrient source, could also contain ligands for both receptors. Further studies are needed to resolve the possible roles of *aliB* and ORF 2 in mediating *S. mitis* serum-associated growth promotion.

**Fig 3:**
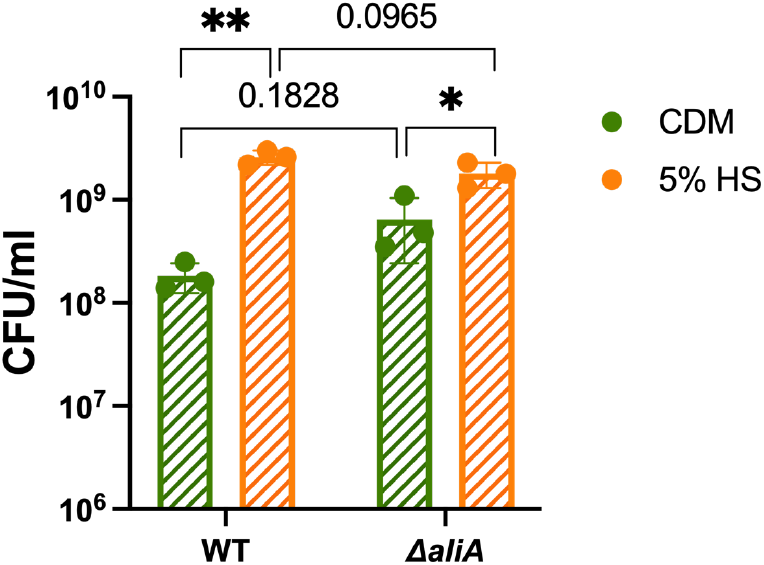
Deletion of *aliA* does not abolish the growth-promoting effects of human serum. Overnight cultures of SM61WT and *ΔaliA* were sub-cultured to 0.1 of OD_600nm_ with fresh CDM with and without supplement of 5% (v/v) human serum (HS). Cell concentrations (CFU/ml) were determined after 8 h of incubation. Three biologically independent replicates were obtained for each tested condition. Statistical analyses were performed with T-test, with significant difference determined by *P*-value < 0.05. Nonsignificant *P*-values are labeled within graph; “*” indicates 0.01 < *P*-value < 0.05; “**” indicate 0.001 < *P*-value < 0.01.

## Conclusions

In this study, we explored the effects of serum supplementation on growth of *S. mitis* and a range of other bacteria that cause bloodstream infections and IE. We observed substantial growth promotion of some, but not all, species and strains tested in the presence of serum. Studies that analyze a larger strain cohort and identify conserved genetic features in strains exhibiting enhanced growth in serum (for e.g. with a genome-wide association study) could reveal a mechanistic basis for this variability. It would also be useful to query multiple strains of *S. pneumoniae, S. pyogenes, S. agalactiae*, and other bacteria used in this study to determine whether variability is also observed. A weakness of our study is the limited number of isolates used for these species.

The mechanism(s) for the observed growth promotion were not elucidated in this study. According to the transcriptomic results, *S. mitis* encounters an environment with more abundant nutrient sources upon the addition of serum, and particularly favors utilization of exogenous lipids rather than *de novo* biosynthesis. The specific serum factors that promote growth of *S. mitis*, which seem to be heat liable, remain uncharacterized, including whether growth promotion is conferred by a single factor or multiple factors. The major heat liable component of serum is the complement, which has antimicrobial activities (59). Though MGS strains have been reported with different complement binding properties (60), complement-induced growth has not been reported before. Moreover, the known virulence factor that confers resistance to complement, PspC (61), is downregulated in this data set. Whether complement contributes to promotion of *S. mitis* growth requires further experimentation. Previously known serum factors that can promote bacterial growth are catecholamines (including dopamine, adrenaline, etc.) and hemoglobin, which are mainly utilized by bacteria as iron sources (62–64). However, no homologs to known transporters that uptake those factors as iron sources have been identified in *S. mitis*, and neither were any iron utilization genes differentially regulated in our data set (Table S1). Finally, it is important to note that CDM, while defined, is also chemically rich, providing amino acids, vitamins and other co-factors, and glucose, among other components. *In situ, S. mitis* would harvest many of these components from the host, or synthesize them. An informative experiment moving forward could be to compare *S. mitis* gene expression in buffer with serum or heat-inactivated serum, which could better illuminate *S. mitis* metabolic processes in the two conditions.

## Materials and methods

### Bacterial strains and culture conditions

All bacterial strains were grown at 37°C, with *Streptococcus* strains grown with 5% CO_2_ supplementation. Unless otherwise stated, *Streptococcus* strains were grown in Todd Hewitt medium (TH medium; BD Biosciences), with *S. pneumoniae* and *S. pyogenes* strains grown in TH medium supplemented with 0.5% yeast extract. *E. coli* were grown in Luria-Bertani (LB) medium. *Enterococcus* strains were grown in Brain-Heart Infusion (BHI) medium. Chemical defined medium (CDM) was made as previously described, with the addition of 0.5 mM choline.(65) Where noted, complete human serum (Sigma Aldrich, H6914) or heat-inactivated human serum (Sigma Aldrich, H5667; treated with 56 ± 2°C for 1 hour by the manufacturer) was added to a final concentration of 5% (v/v), and ZnCl_2_ was added to a final concentration of 0.25 mM. Viable cell concentrations were determined by serial dilution with phosphate buffered saline (PBS) and quantification of colony forming units (CFUs). All bacterial strains and plasmids used in this research are shown in Table 3. All primers used in this research are listed in Table 4.

**Table 4:**
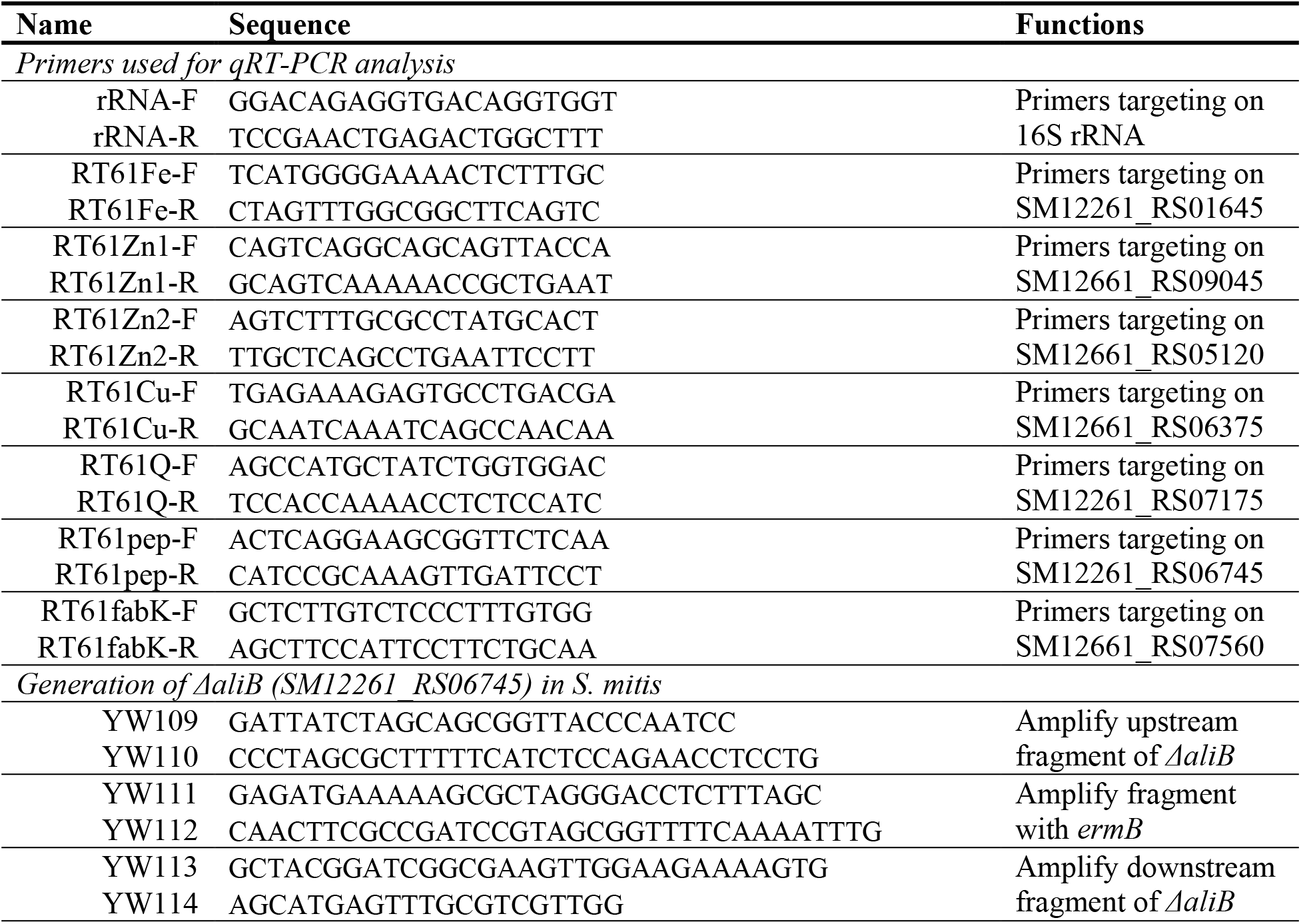
Primers used in this research.

### RNA sequencing

Single colonies of *S. mitis* ATCC 49456 (SM61) were grown in CDM overnight followed by dilution to an OD_600nm_ of 0.1 with fresh pre-warmed medium into two replicates. When the OD_600nm_ values of the diluted cultures reached 0.3, complete human serum was added to one set of the cultures to a final concentration of 5% (v/v), followed by another 30 min of incubation at 37°C with 5% CO_2_. Then the cells were pelleted, resuspended into 450 μl media, and mixed with 900 μl RNAProtect (Qiagen) for stabilizing RNA. After incubation at room temperature for 10 minutes, cells were pelleted at 5000 x g for 10 minutes and stored at -80°C prior to further processing.

Total RNA was extracted with the following procedure. Cell pellets were resuspended into 1 ml of ice-cold RNABee (Tel-Test, Inc.), transferred into Lysine Matrix B tubes (MP Biomedicals), followed by 6 cycles of bead-beating at 6.5 m/s for 45 seconds, with 5 minutes on ice between cycles (FastPrep-24™ MP Biomedicals). Then, 250 μl chloroform was added to each of the samples, followed with ∼ 30 seconds shaking by hand followed by incubation on ice for 10 minutes. After centrifugation at 17000 g for 15 minutes, the aqueous layer was removed and mixed with an equal volume of isopropanol, followed by incubation on ice for >1 hour. RNA pellets were obtained via centrifugation at 17000 g for 15 minutes at 4°C, followed by two washes with ice-cold 70% ethanol. The RNA pellets were air-dried before rehydration with RNase-free water.

Removal of DNA was conducted with Recombinant DNase I (Roche Diagnostics) per the manufacturer’s instructions, followed by cleaning with the GeneJET RNA Cleanup and Concentration Micro Kit (Thermo Scientific). Lack of DNA contamination was confirmed with PCR using primers rRNA-F and rRNA-R (Table 4). RNA sequencing of samples with RNA integrity number (RIN) above 9.4 were performed by the Genome Center at The University of Texas at Dallas (Richardson, TX). Specifically, rRNA was removed with RiboMinus kit (Invitrogen), sequencing libraries were prepared with TruSeq Stranded mRNA Library kits (Illumina) and sequenced with Illumina Nextseq 500.

Analysis of the RNA sequencing results was performed with the CLC Genomics Workbench (version 20; Qiagen). The *S. mitis* ATCC 49456 genome sequence (NZ_CP028414.1) was used as reference. Sequence reads that aligned to rRNA genes were removed from further analysis. Reads per kilobase of transcript, per million mapped reads (RPKM) values of each annotated gene was calculated and used for analysis of differential expression at the transcript level. Validity of replicates were confirmed with principal component analysis (Fig S2) and scatter plot with linear regression using the RPKM values (Fig S3), R^2^-values of the comparisons between replicates under the same testing condition are all above 0.9. RNA sequencing data for all ORFs are shown in Table S1. Differentially expressed genes were determined by absolute fold change > 1.5, false discovery rate (FDR) *P*-value < 0.01, and maximum RPKM value > 10.

### Quantitative reverse transcriptase PCR (qRT-PCR)

Primers used for qRT-PCR were designed via the online program Primer3 with 60±2°C set as the optimized annealing temperature (66). Specificity of the primers was confirmed with PCR using purified genomic DNA as template and analysis of products by agarose gel electrophoresis. cDNA was generated from total RNA samples described above using SuperScript3 (ThermoFisher) per the manufacturer’s instructions and using 1 μg total RNA as template, followed by cleaning with the QIAquick PCR purification Kit (Qiagen). cDNA concentration was measured by Nanodrop (Thermo Scientific) and diluted to 100 ng/μl for use in qRT-PCR. qRT-PCR reactions were prepared with AzuraQuantTM Green Fast qPCR Mix LoRox (Azura Genomics) per the manufacturer’s instructions, using 1 μl cDNA per reaction for the detection of 16S rRNA and 5 μl cDNA per reaction for the detection of target genes. qRT-PCR reactions were performed on the MyGo Pro machine (Azura Genomics) and analyzed with the MyGo Pro PCR Software (v3.2, Azura Genomics). Relative transcript levels of the target genes were calculated with the ΔΔCt method with normalization to the Ct numbers of 16S rRNA. For each sample, at least three biological independent replicates were obtained. Statistical analysis was analyzed with one-way ANOVA, and significant differences were determined by *P*-value < 0.05.

### Gene ortholog identification

Due to the close evolutionary relationships and genomic similarity between *S. mitis* and *S. pneumoniae*, as they share > 900 core genes (17, 67), previous studies of *S. pneumoniae* regulatory systems were used as the main reference for the interpretation of the putative regulatory relationships among the differentially expressed genes. Specifically, gene orthologs were identified through using the BLASTp function against the NCBI database (68). Proteins of interest from *S. pneumoniae* were used as input to query the nonredundant protein database of *S. mitis* ATCC 49456^T^ (taxid: 246201). Orthologs were determined by the lowest E value (< 10^−88^) of each inquiry with a query coverage of > 94%.

### Mutant generation

*S. mitis* mutants are generated as described before (69). Specifically, overlapping PCR was used to sequentially join a 2 kb fragment from upstream of the target gene, a 1 kb fragment containing *ermB* in the reverse orientation, and a 2 kb fragment from downstream of the target gene. The purified 5 kb fragment was transformed into *S. mitis* competent cells as previously described (69). Mutant candidates were selected on media with 20 μg/ml erythromycin and confirmed with Sanger sequencing (MGH CCIB DNA Core, Cambridge, MA).

### Accession numbers

Raw sequencing data was submitted to the NCBI database with BioProject ID of PRJNA784985 and BioSample accessions of SAMN23523627-32.

## Acknowledgements

We gratefully thank the Genome Center at The University of Texas at Dallas for the services to support our research. This research is supported by grant R01AI148366 from the National Institutes of Health and the Cecil H. and Ida Green Chair in Systems Biology Science to K.L.P.

## Supplementary information

In excel file:

**Table S1:** Complete sequencing analyses data

**Table S2:** Homologs to *S. pneumoniae* glutamine transporters

**Fig S1:**
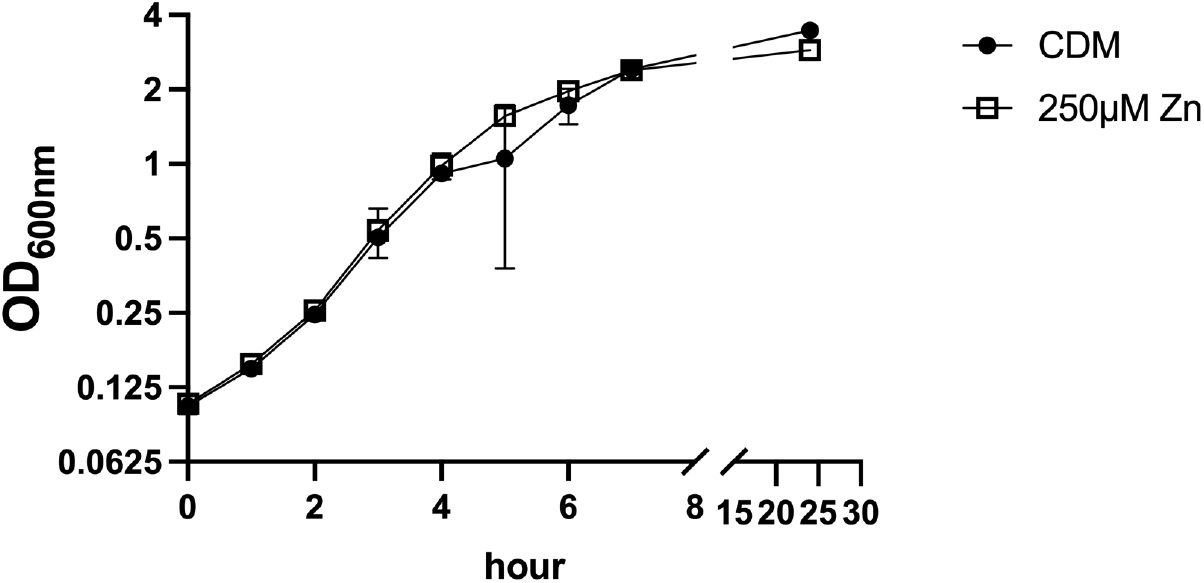
Growth curve of *S. mitis* ATCC 49456^T^ (SM61) grown in either chemically defined medium (CDM) or CDM with 250 μM ZnCl_2_. Two biologically independent replicates were obtained for each tested condition.

**Fig S2:**
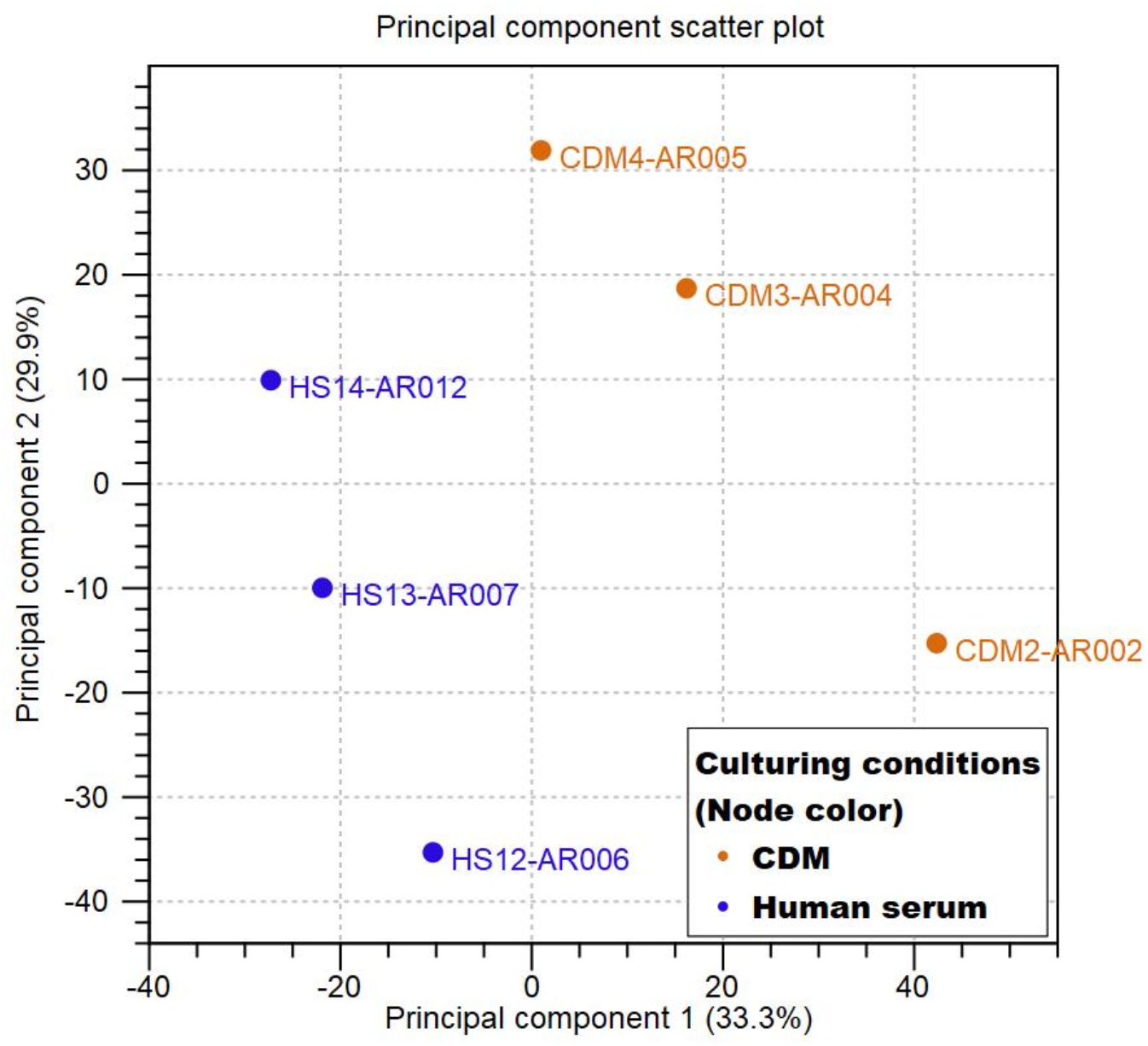
Plot of the principal component analysis of the RNA sequencing samples. Samples obtained from plain chemically defined medium (CDM) are represented by orange dots; while samples obtained from serum supplemented cultures are represented by blue dots. Plot figure is generated through the PCA function of the CLC Genomic Workbench.

**Fig S3:**
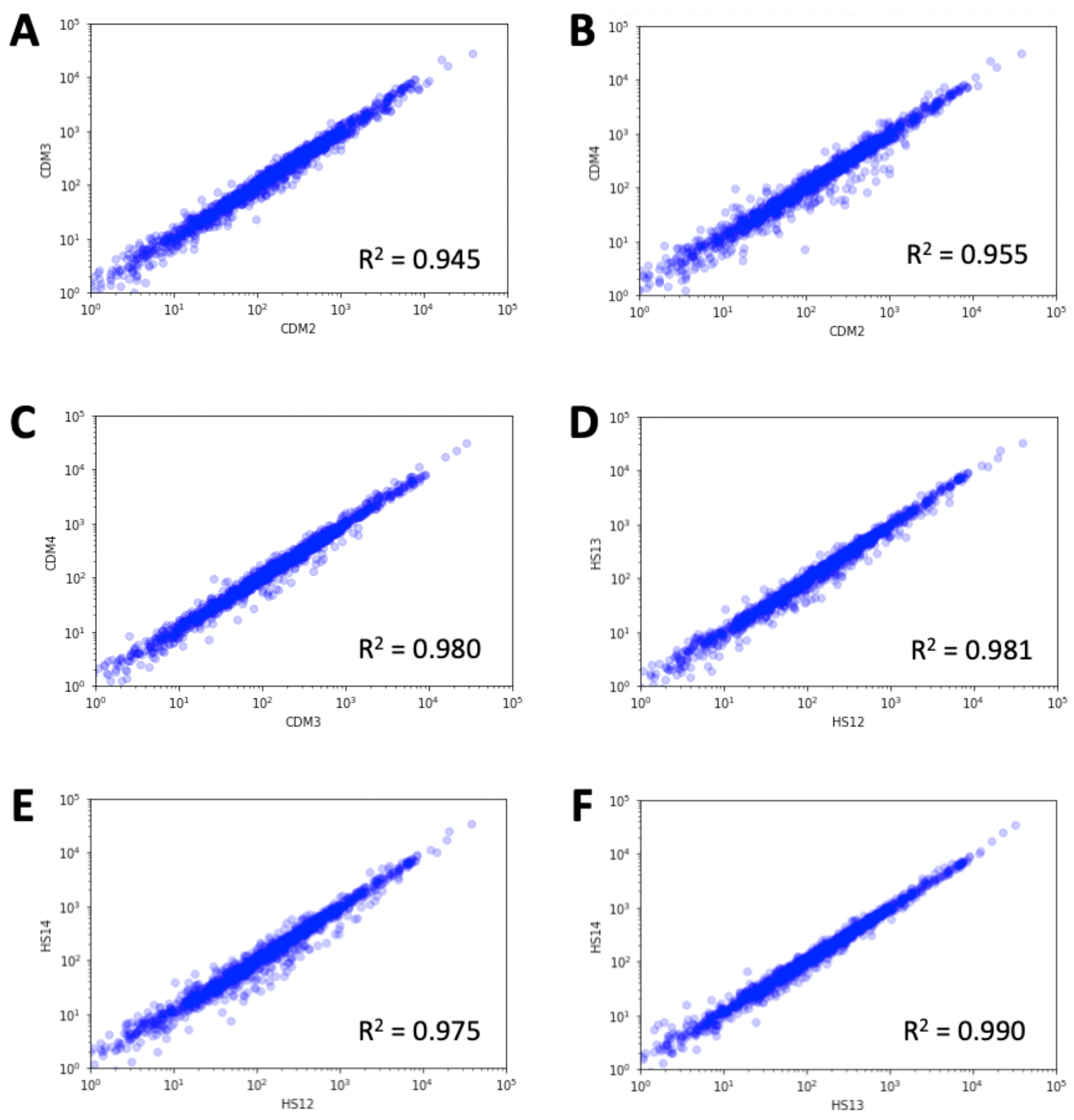
Replicate validation of the RNA sequencing samples via scatter plot analyses. Axis labels indicate the samples in comparison. “CDM” stands for samples grown in chemically defined medium. “HS” stands for samples grown in medium supplemented with human serum to a final concentration of 5% (v/v). R^2^-values are labeled within each plot.

